# Cell cycle dynamics during diapause entry and exit in an annual killifish revealed by FUCCI technology

**DOI:** 10.1101/522417

**Authors:** Luca Dolfi, Roberto Ripa, Adam Antebi, Dario Riccardo Valenzano, Alessandro Cellerino

## Abstract

**Background:** Annual killifishes are adapted to surviving and reproducing over alternating dry and wet seasons. During the dry season, all adults die and desiccation-resistant embryos remain encased in dry mud for months or years in a state of quiescence, delaying hatching until their habitats are flooded again. Embryonic development of annual killifishes deviates from canonical teleost development. Epiblast cells disperse during epiboly, and a “dispersed phase” precedes gastrulation. In addition, annual fish have the ability to enter diapause and block embryonic development at the dispersed phase (diapause I), mid-somitogenesis (diapause II) and the final phase of development (diapause III).

Developmental transitions associated with diapause entry and exit can be linked with cell cycle events. Here we set to image this transitions in living embryos.

**Results:** To visibly explore cell cycle dynamics during killifish development in depth, we created a stable transgenic line in *Nothobranchius furzeri* that expresses two fluorescent reporters, one for the G_1_ phase and one for the S/G_2_ phases of the cell cycle, respectively (fluorescent ubiquitination based cell cycle indicator, FUCCI). Using this tool, we observed that, during epiboly, epiblast cells progressively become quiescent and exit the cell cycle. All embryos transit through a phase where dispersed cells migrate, without showing any mitotic activity, possibly blocked in the M phase (diapause I).

Thereafter, exit from diapause I is synchronous and cells enter directly into the S phase without transiting through G_1_. The developmental trajectories of embryos entering diapause and of those that continue to develop are different. In particular, embryos entering diapause have reduced growth along the medio-lateral axis. Finally, exit from diapause II is synchronous for all cells and is characterized by a burst of mitotic activity and growth along the medio-lateral axis such that, by the end of this phase, the morphology of the embryos is identical to that of direct-developing embryos.

**Conclusions:** Our study reveals surprising levels of coordination of cellular dynamics during diapause and provides a reference framework for further developmental analyses of this remarkable developmental quiescent state.

**List of Abbreviations:** In this paper, we will refer to several developmental stages or morphological structures using abbreviations. To make the reading easier, we resume here a list of all the abbreviations, to which the reader can refer at any time.

WS: Wourms Stage. Developmental stage referring to the embryonic description made by Wourms for the killifish species A*ustrofundulus limneus*.
YSL: Yolk syncytial layer. A layer of cells that form a syncytium and that are in direct contact with the yolk. This is the most internal layer, through this layer nutrients from the yolk can be delivered to the upper layers.
EL: Epiblast layer: A layer of cells composed by blastomeres that divides actively during development and will take part in the generation of the several embryonic and fish major structures like head tail trunk and organs.
EVL: Enveloping layer. A thin layer of cells that envelopes all the embryo. It is the most external layer. The cells belonging to this layer are big with big nuclei that do not divide.
DI: Diapause I. A dormancy stage peculiar of annual killifish species that occurs after the completion of epiboly, during the dispersed phase.
DII: Diapause 2. The second and most important dormancy stage of annual killifish species. Fish can stop in DII only entering a different developmental trajectory after the reaggregation phase. The final developmental block occurs at the mid somitogenesis stage.
DC: Diapause Committed embryo. An embryo that undertook the Diapause II trajectory of development and that will stop for sure in Diapause II during the somitogenesis stage.
DD: Direct Developing embryo. An embryo that is following the not diapause II developmental trajectory. These embryos grow more in lateral size during somitogenesis and never stop their development in this phase.

## Introduction

Annual killifishes inhabit temporary habitats that are subject to periodic desiccations [1]. In order to survive these extreme conditions, their eggs are laid in the soft substrate and remain encased in the dry mud where they are relatively protected from desiccation and can survive for prolonged periods during the dry season and regulate their development in anticipation of the ensuing rainy season. When their habitats are flooded, these embryos hatch, develop rapidly and spawn the next generation before water evaporates [2-6]. This seasonal life cycle comprising embryonic arrest is widespread in arthropods from temperate climates, but it is unique among vertebrates.

As an adaptation to seasonal water availability, embryonic development of annual killifishes deviates from canonical teleost development for three main distinctive traits. The first is a slow cell cycle during early cleavage. While embryos of non-annual teleost fishes execute one cell division every 15-30 minutes during the first divisions after fertilization, the rate of early cell division in annual killifishes can reach almost two hours [7]. As a result, an annual killifish embryo can be still in the blastula stage while a non-annual killifish embryo fertilized at the same time has started somitogenesis.

The second trait is the dispersion of epiblast cells during epiboly and a decoupling between epiboly and gastrulation. When epiboly starts, the epiblast cells delaminate, assume an amoeboid shape and migrate towards the other pole of the egg. This migration is physically guided by the spreading of the extra embryonic enveloping layer [8]. In annual killifishes, the embryo at the end of epiboly consists only of extraembryonic structures and separated epiblast cells that migrate randomly over the yolk surface in a unique developmental stage named dispersed phase [6]. The dispersed phase can last for several days and the embryonic axis is formed by migration of the epiblast cells towards a point where they reaggregate and form the embryonic primordium. This peculiar stage is named reaggregation phase [6]. In several teleosts, including zebrafish, gastrulation and axis formation take place during epiboly. However, in annual killifishes the formation of the three embryonic layers, which happens during gastrulation, takes place after epiboly during the late aggregation phase as demonstrated by live cell imaging and by the expression of the blastopore markers *goosecoid* and *brachyury* [9].

The third unique feature of annual killifish development is the ability to enter diapause. Diapause is a state of dormancy that retards or blocks embryonic development in anticipation of predictable cyclic hostile conditions. Diapause is widespread among arthropods from temperate climates that spend in diapause the coldest part of the year. Annual killifish embryo can arrest in diapause in three specific phases of development: during the dispersed phase (diapause I), at mid-somitogenesis when most organs are formed (diapause II) or at the final stage of development (diapause III) [4]. Duration of diapause is highly variable and diapauses are not obligatory [4]. These adaptations are interpreted as bet-hedging strategies that ensure survival in an unpredictable environment, which are typical of seed banks [3, 10]. Under appropriate conditions, such as high temperature or under the influence of maternal factors, embryos can greatly shorten and possibly skip all three diapauses and proceed through direct development [11-14]. Diapause II is not a simply a phase of developmental arrest, but direct development and diapause are alternative developmental trajectories, characterized by different morphologies. In particular, during somitogenesis, the embryos committed to diapause II grow in the longitudinal direction but are impaired in transversal growth and therefore have reduced transversal diameter of head and body as compared to direct development embryos [13, 15]. This difference is detectable already at the start of somitogenesis and it is observed in multiple independent clades that have evolved diapause [15] and can have an impact on post-hatch life history traits [10]. Diapause II can be extremely prolonged, and in some species can last for years (personal observation, AC, DRV).

One species of annual killifish has recently become a relatively widespread experimental model: *Nothobranchius furzeri*. This is the shortest-lived vertebrate that can be cultured in captivity and it replicates many typical phenotypes of vertebrate and human aging [2, 16-18]. For this reason, it has been used as an experimental model to investigate the effects of several experimental manipulations on aging [10, 19-25]. Natural habitats of this species can last as short as a month [26, 27] and yet are able to sustain a viable population since sexual maturity can be reached within two weeks from hatching [28]. *N. furzeri* therefore represent an extreme case of compressed lifespan among annual killifishes and in natural conditions it spends longer time as embryos in diapause than in post-hatch stages.

Molecular studies of killifish embryonic development are scarce. It is known that diapause II is characterized by reduced protein synthesis, cell cycle arrest and remodelling of mitochondrial physiology and it is controlled by insulin-like growth factor 1 signalling [29-35]. Recently, RNA-seq studies have shown that, gene-expression patterns during diapause resemble those observed during aging [36] and vitamin D signalling controls to the choice between direct development and diapause II [37]. While early embryonic development following fertilisation have been in part studied [7-9], the characterization of the physiological and molecular events occurring during diapause I has been started to be investigated only recently [35, 38, 39]. Here, to investigate entry and exit form diapause I and II, we created a stable transgenic line in *N. furzeri* that expresses fluorescent reporters for cell cycle phases [40]. This tool enabled us to shed light on the cell cycle characteristics of cells during diapause I and to demonstrate that diapause exit is characterized by rapid and synchronous cell cycle reactivation. These characteristics appear unique among vertebrate embryos.

## Results and Discussion

To identify a suitable promoter for FUCCI reporter lines, we tested the activity in *N. furzeri* of the zebrafish ubiquitin (ubi) promoter [41] using EGFP as reporter. We observed ubiquitous expression of the EGFP from the second day of development into adulthood in this Tg *ubi:egfp* line (Figure 1D).

**Figure 1:**
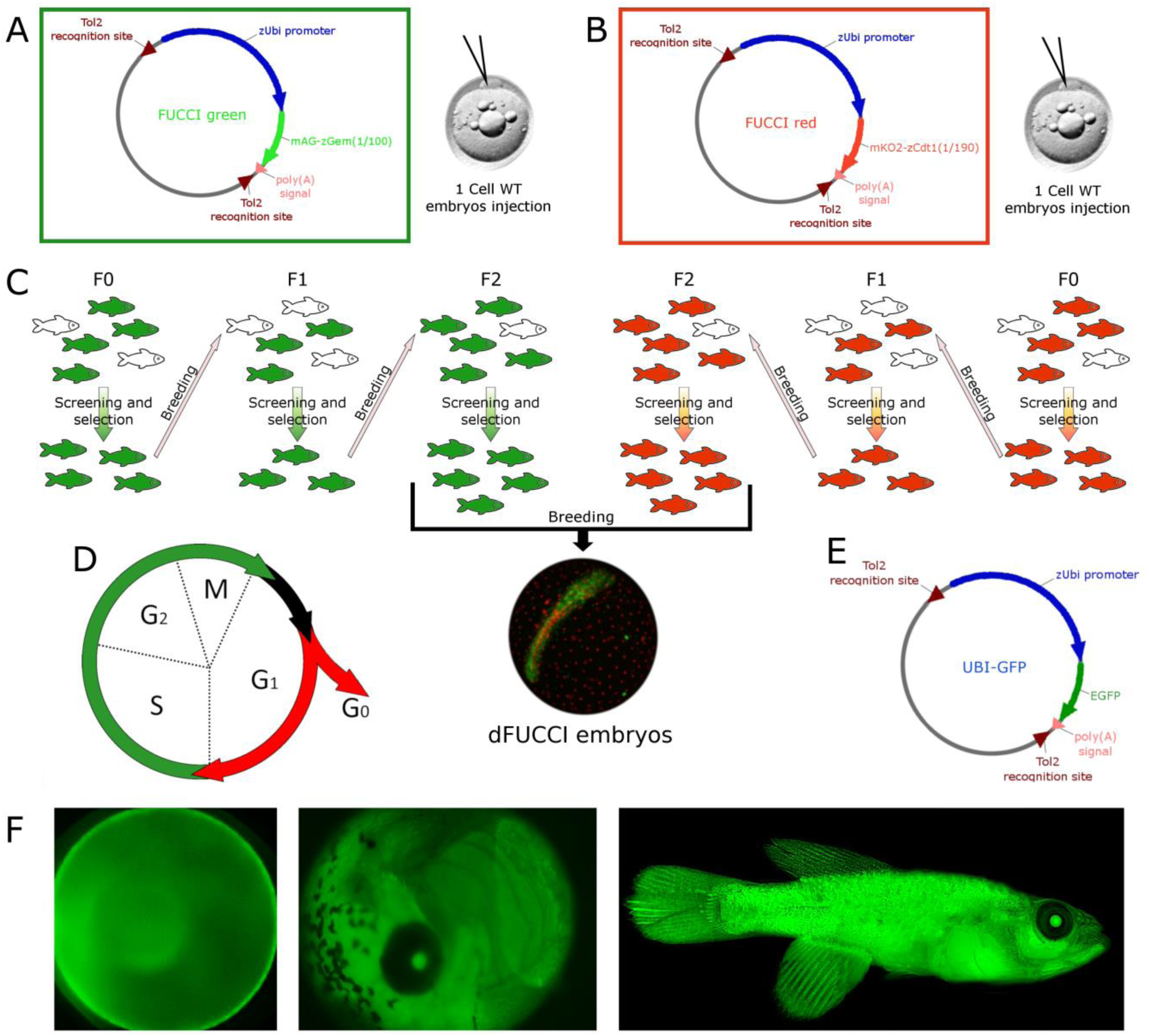
FUCCI transgenic line generation. A and B are schematic representations of FUCCI green and red constructs, respectively. FUCCI constructs were injected separately in different 1-cell stage fertilized eggs. Positive eggs were raised into adult fish, bred and screened for 3 generations (C). F2 FUCCI green fish were finally bred with F2 FUCCI red fish to generate double FUCCI embryos, which were used for most experiments (C). D is a schematic representation of how FUCCI technology works, cells are green during S/G_2_/M phases, colorless between M and G_1_ and red in G_1_ and G_0_ phases. E is a schematic representation of zUbiquitin-EGFP construct. EGFP expression in Nothobranchius embryos and adult fish is shown (F)

Two different FUCCI transgenic lines [40] were successfully generated using the Tol2 transgenesis system [42-44],: i) a “FUCCI green” reporter (Azami green - Geminin), which is activated during the S/G2/M phase of the cell cycle [40] and ii) a “FUCCI red” reporter (Kusabira orange - Cdt1), which is activated during the G1 phase of the cell cycle [40]. Both transgenes were placed under the control of the zebrafish *ubi* promoter (Figure 1 A,B).

Adult F0 transgenic fish were screened for fluorescence and bred one to another. F1 fish showing the expected fluorescence pattern were interbred in order to increase the number copies of FUCCI reporter cassettes in their genome, thereby enhancing the fluorescence signal in the F2 generation (Figure 1C). F2 transgenic fish were used to characterize the expression pattern of the FUCCI reporters at different developmental stages.

We describe the expression pattern of the transgenes in two parts: the first part provides a detailed description of the fluorescent signal in the single lines (FUCCI green and FUCCI red, respectively) during focal stages of embryonic development and in adult life. The second part is focused on the double transgenic line, where both the red and green signals are present. In this part, changes in relative intensity of the two signals are described and interpreted with respect to the dynamic processes that characterize the different phases of killifish embryonic development.

### Part 1

#### FUCCI red

The FUCCI red signal localized in the nuclei of cells in every stage of *N. furzeri* life. During the dispersed phase (Figure 2A,B), two cell types expressed red fluorescence: large cells of the enveloping layer (EVL) and some smaller cells of the epiblast (Figure 2C, arrows). The nuclei of EVL cells ranged from 22-27 um in diameter and formed a regular array over the yolk surface. The nuclear diameter of epiblast cells was smaller, on the order of 7-9 um. Both these cell types were red, but the epiblast cells were most likely in G1, since their fluorescence faded and increased over time course of hours, indicating that they were engaged in the cell cycle. By contrast, the EVL cells appeared to be in G0, since the red fluorescence never faded, and lasted throughout embryonic development, until hatching.

**Figure 2:**
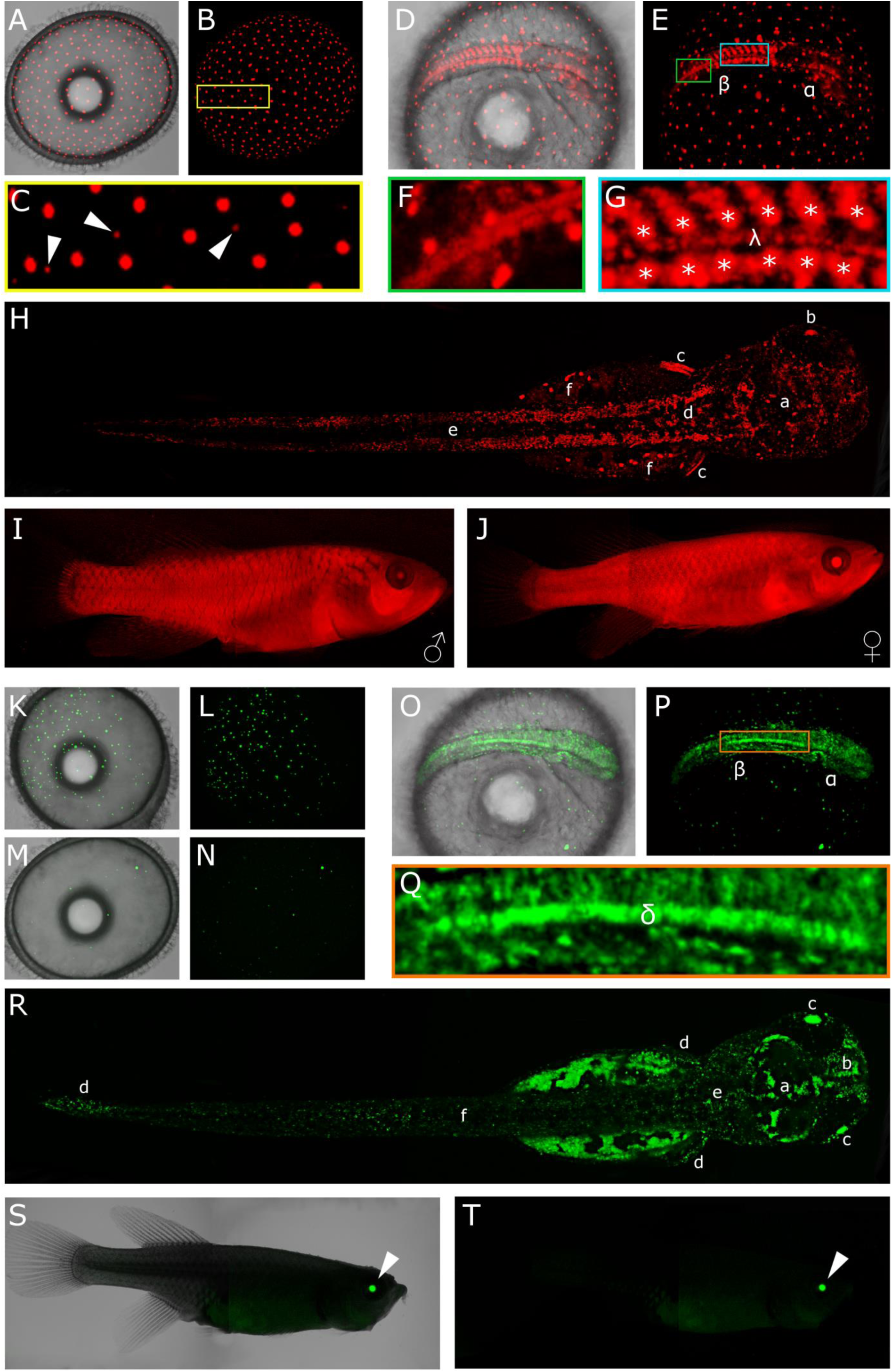
FUCCI green and FUCCI red characterization. A to J show FUCCI red expression at different developmental stages. K to T show FUCCI green expression at different developmental stages. A,B,C,K,L,M,N dispersed phase. D,E,F,G,O,P,Q somitogenesis stage. H,R hatched fry. I,J,S,T adult fish.

At the somitogenesis stage (Figure 2D,E), the regular distribution of the EVL cells remained unchanged. All the other red cells in the embryo showed a patterned distribution that was particularly striking in the embryo trunk, where the somites were clearly delineated. In the older more rostral somites (Figure 2G), the inner part of the somites showed a high concentration of red cells, while in the more caudal part of the embryo (Figure 2F) – where somites were still forming – the red cells were more spread and diffused. Once the new somite pair was completely formed, red cells increased in numbers, becoming more dense and localized in the inner part (Figure 2G, stars), becoming constitutively red. Just below the notochord midline, another narrow streak of red cells extended from the tip of the head region to the tip of the tail (Figure 2G, λ). Also the head region contained several areas with red cells, but these were quite rare, spread out and did not demarcate specific areas (Figure 2E, α).

Hatched fry had large fraction of red cells (Figure 2H). The lateral muscles of the trunk (Figure 2H, d) and of the tail (Figure 2H, e) harboured a large number of red cells. In the head region nearly every part of the brain had some spread red cells or red cells aggregates (Figure 2H, a). The lens (Figure 2H, b) showed strong red fluorescence at this stage. This could be an artefact due to the high protein stability in this region and lack of degradation of the FUCCI reporter. What remained of the yolk at this stage was still surrounded by large red cells belonging to the EVL, blocked in G0 and not cycling (Figure 2H, f). Lastly, a large patch of red cells was observed corresponding to the pectoral fins (Figure 2H, c).

The adult FUCCI red transgenic fish appeared completely red under fluorescence since many cells were in G0 or possibly G1 phase (Figure 2I,J). Males and females showed a pattern that was virtually identical, and the signal intensity was comparable between different specimens (Figure 2I,J).

#### FUCCI green

During the dispersed phase, different proportions of green epiblast cells were detected in FUCCI green embryos. The number of green nuclei observed varied between 20 (Figure 2M,N) to 200 (Figure 2K,L), with a size range between 7-25 um in diameter, typically showing a higher amount of the smaller cells. During this developmental stage, cells arranged randomly over the yolk surface.

In developing embryos, during somitogenesis proliferating green cells were detected in every part of the forming embryos (Figure 2O,P). The signal was moderately strong in the trunk, in the tail (Figure 2P, β) and in the head primordia (Figure 2P, α). The maximum intensity of the signal, the maximum density of proliferating cells, was limited to a narrow region along the midline, that extended from the end of the head to the end of the tail, between the yolk surface and the lowest part of the somites (Figure 2Q, orange box, δ). Many green cells migrated over the yolk surface during all of somitogenesis.

In the hatched fry, the proliferative regions in the embryos were more defined. In the torso and the tail, green cells were spread out but homogeneously distributed (Figure 2R, f). Slightly more dense green cells proliferated in the caudal and pectoral fins (Figure 2R, d) and at the base of the head, in the hindbrain (Figure 2R, e). The forebrain (Figure 2R, a) showed a clear pattern with thick and dense aggregates of green cells at the borders, in the proliferating niches, with almost no green cells in the inner part, appearing completely dark. The olfactory bulb (Figure 2R, b) in the fry were one of the major proliferating regions, composed by thick streaks of green cells. Lastly, the lens appeared green (Figure 2R, c), but as for the FUCCI red transgenic line, this could be an artefact due to lack of degradation of the FUCCI reporter. In adults, no green cells could be detected with a stereomicroscope and the only green signal detectable was confined to the region of the eyes (Figure 2S,T, arrow).

### Part 2

F2 fish were crossed (FUCCI red with FUCCI green), generating double FUCCI green/red embryos (dFUCCI), which were analysed by means of time-lapse confocal imaging. Embryos from this cross were imaged for periods spanning from hours to days at different stages of development from the end of epiboly to late somitogenesis, i.e. past the stage when embryos entered diapause II. The stacks of images of each time point were then processed with IMARIS software to perform particle tracking and counting.

These experiments required occupancy of the set-up for considerable amounts of time, as every single time lapse acquisition lasted from 8 hours to 4 days, limiting the number of replicates available for each stage (Table 1). This study was designed to provide an overview, as complete as possible, of *N. furzeri* embryonic development (compromising on the number of replicates) as opposed to deep analysis of only one specific stage (e.g. reaggregation) with a larger number of replicates. Because the morphology of Southamerican and African annual killifishes are comparable, we follow here the staging developed by Wourms for the South American annual killifish *Austrofundulus limnaeus* [6] for reference.

**Table 1:** Overview of dFUCCI embryos imaged for each developmental stage. The table shows the number of embryos acquired at each developmental stage. From stage to stage embryos acquired could be the same, acquired progressively during its development, or different ones. WS means Wourms stage [12].

| Embryonic Stage |  | Amount of embryos imaged |
| --- | --- | --- |
| Dispersed Phase (WS 19-20) | Early (small number of green cells) | 4 |
|  | Late (larger number of green cells) | 6 |
| Dispersed Phase transition (from few to many green cells) |  | 4 |
| Reaggregation Phase (WS 21-25) |  | 4 |
| Extension Phase (WS 26) |  | 4 |
| Somitogenesis (without diapause II arrest) (WS 29-33) |  | 6 |
| Diapause II arrest and release |  | 4 |

In his work, Wourms described the stages of killifish development from egg fertilization to fry. He dived the development in a total of 46 stages, 43 pre-hatching and 3 post-hatching. Stage 1 defines the freshly fertilized 1 cell stage embryo while stage 46 describes the fry after digestion of the yolk that is starting to hunt prey..

In our work, we focus on a subset of these stages and, more precisely, we describe development from stage 19 to 33, which correspond to the phase between completion of epiboly and beginning of the dispersed phase to an advanced stage of somitogenesis.

### End of epiboly (WS 19)

Detection of fluorescence signal with a confocal microscope was not possible before the stage of 70% epiboly because the expression of the fluorescent reporters was too weak: earlier stages of epiboly were described previously by injecting synthetic RNA coding for FUCCI reporters [7]. During epiboly, three cell layers become defined: the yolk syncytial layer (YSL), the enveloping layer (EVL) and the epiblast layer (EL) (Figure 3).

**Figure 3:**
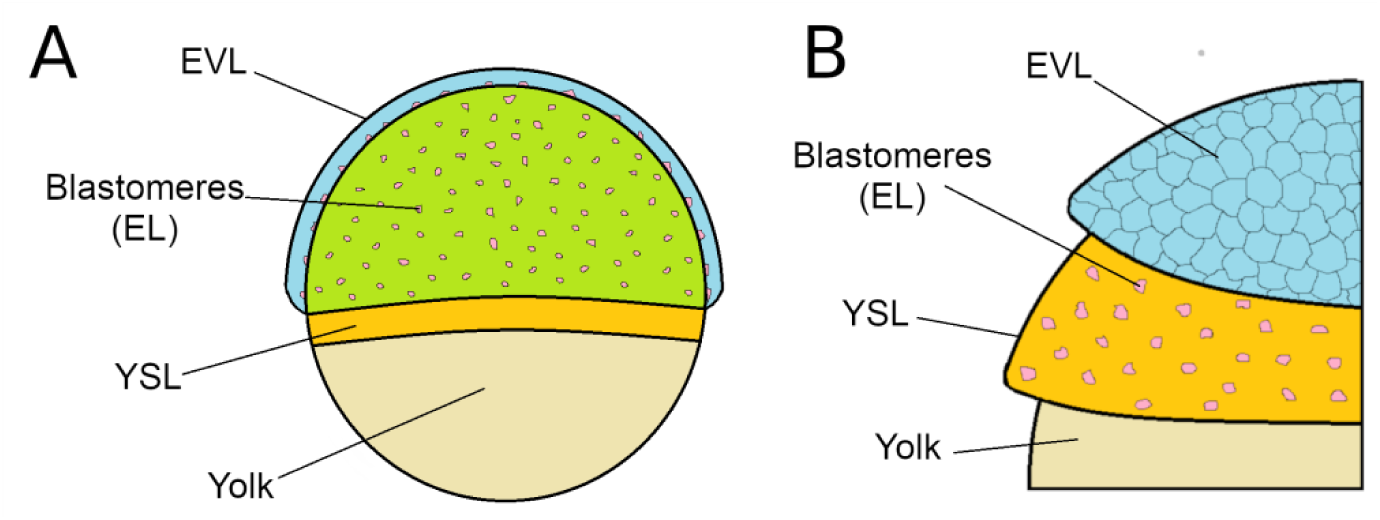
Schematic representation of embryonic layers in annual killifish species. Yolk syncytial layer (YSL) is made of cell forming a syncytium and in direct contact with the yolk. On top of this layer migrate and divide the blastomeres, a discrete population of cell composing the epiblast layer. The enveloping layer (EVL) is the layer of cells on top, a thin layer that completely envelope the other two. Two representations are shown, a lateral semi-transparent view (A) and a cutaway representation (B).

Two of these three cell types could be clearly distinguished based on dFUCCI signal: the EVL cells and the EL cells, which could be distinguished based on their non-overlapping spans of nuclear sizes (21-27 μm vs. 8-12 μm, respectively). The EL cells (showing either green or red fluorescence) migrated in an apparently random direction in the space between the YSL and the EVL. Remarkably, these cells continued their movements also once epiboly was completed. Random movements of EL cells in the dispersed phase were originally reported by Lesseps et al., in the ‘70s [45] by means of bright-field microscopy and were later confirmed by us [6].

EVL cells maintained red fluorescence for the entire span of subsequent development, possibly indicating arrest in G_0_ phase, which is coherent with the not proliferating status of these cells. Additionally, their number remained stable around 200 in the portion of the embryo that could be imaged (corresponding roughly to the superior pole) (Figure 4). EVL cells showed a directional movement until the completion of epiboly (Wourms stages 18-19) (Movie as additional file1), when they reached their final position and constituted a syncytium, which was then maintained during the ensuing development. This physical movement and positioning of EVL cells plays an important role during the development of killifish embryos. The spreading of EL cells is indeed instructed by mechanical interactions between EVL and EL cells [8].

**Figure 4.**
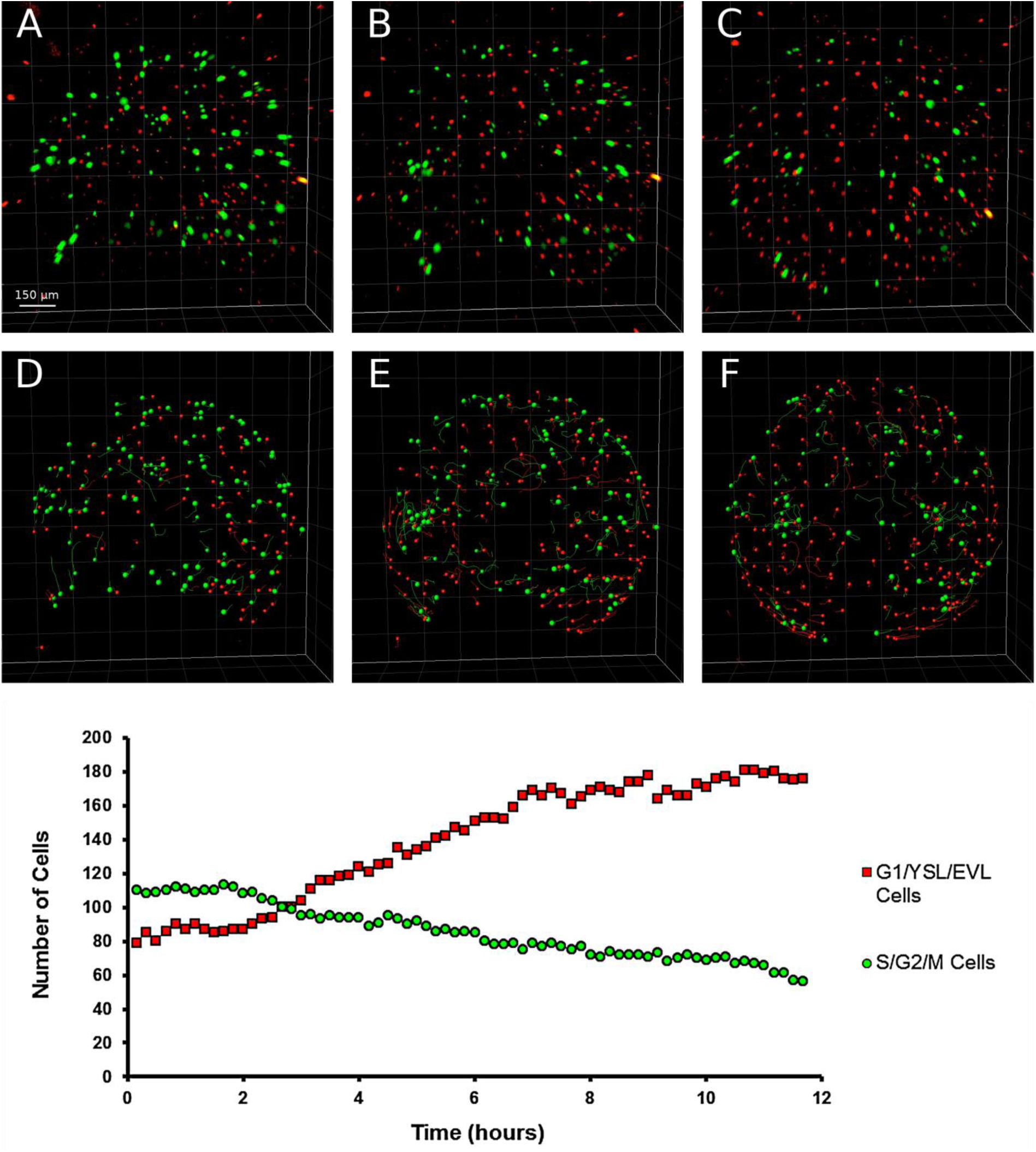
Cell dynamics during epiboly (Wourms stages 18-19). When epiboly is ongoing, both green and red cells are present in dFUCCI embryos (A). As long as epiboly proceeds (B, C) the number of green EL cells gradually and slowly decreases over time while the number of red EVL cells increases until the reaching of a plateau (graph hours 7-12). The cells in the field of view were easily tracked and counted transforming the dots in particles with IMARIS(D,E,F). The images and graph refer to the acquired portion of the embryos, corresponding to the superior hemisphere.

EVL cells formed a defined and regular architecture, tiling the entire surface at an average distance of about 80 μm from each other. Therefore, we used the position of these nuclei as a reference to correct for yolk movements that often occurred in the embryos during development, allowing for precise tracking of the movements of all the other cells and developing structures (Figure 5).

**Figure 5.**
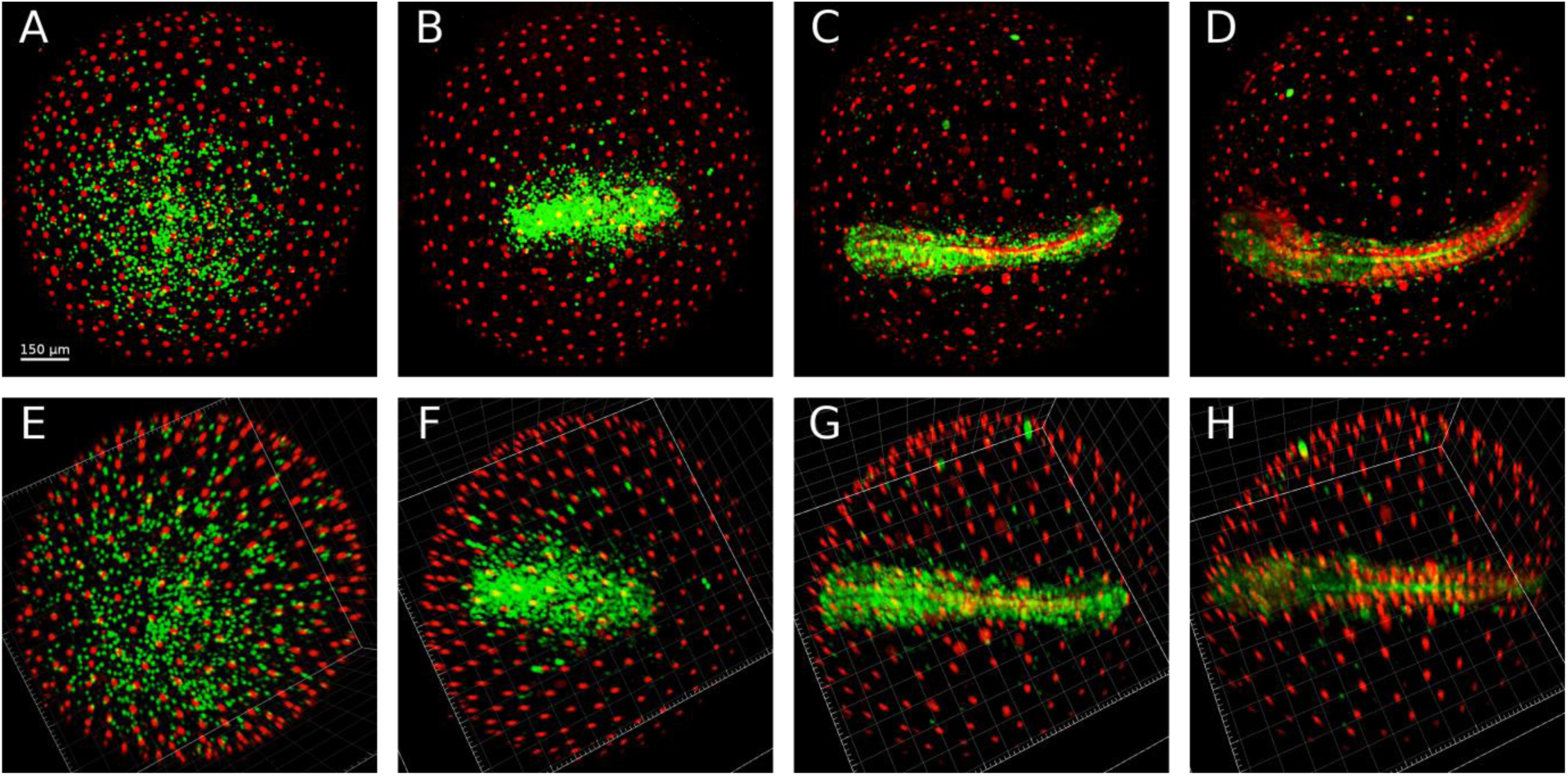
ESL nuclei as reference point for drift correction. Embryos continuously move throughout development (A,B,C,D) but EVL nuclei do not move once they reach their final position at the end of epiboly. These nuclei can therefore be used as reference system and drifts and rotations that occurs during development can be corrected using their position (E,G,F,H). Correcting the drifts allows a more precise and reliable cell tracking and data analysis.

### Early dispersed phase (Wourms stages 19-20) and diapause I

When epiboly ends and the dispersed phase begins, the number of detectable green and red EL cells was reduced from more than 200 to less than 70, in a ratio of almost 1:1 (Figure 4 and 6C, left panel). These cells are actively moving in a seemingly random fashion (Figure 6B and 6C, right panel). It is important to remark that the decline in green and red cells does not correspond to cell loss, as EL cells are clearly detected by bright-field microscopy at this stage (Movie ad additional file 2, min 0.53 to 1.10). Previous studies of early cleavages by means of injection of synthetic FUCCI mRNA [7] also reported a “dark phase” when neither red nor green fluorescence is detectable. The majority of EL cells appear to be locked in this “dark phase” during early dispersed stage. At the moment, we can only speculate as to the cell-cycle characteristic of these “invisible” cells. In mouse cells in culture, FUCCI fluorescence is detectable only during S/G_2_/M or G_1_/G_0_ phases but not at the late stages of M phase, so that a brief period signal absence ensues at the end of the green phase and precedes cytokinesis [46]. So, it is likely that during the early dispersed phase cells are locked in telophase and cannot proceed to cytokinesis. However, we lack direct support for this hypothesis.

**Figure 6.**
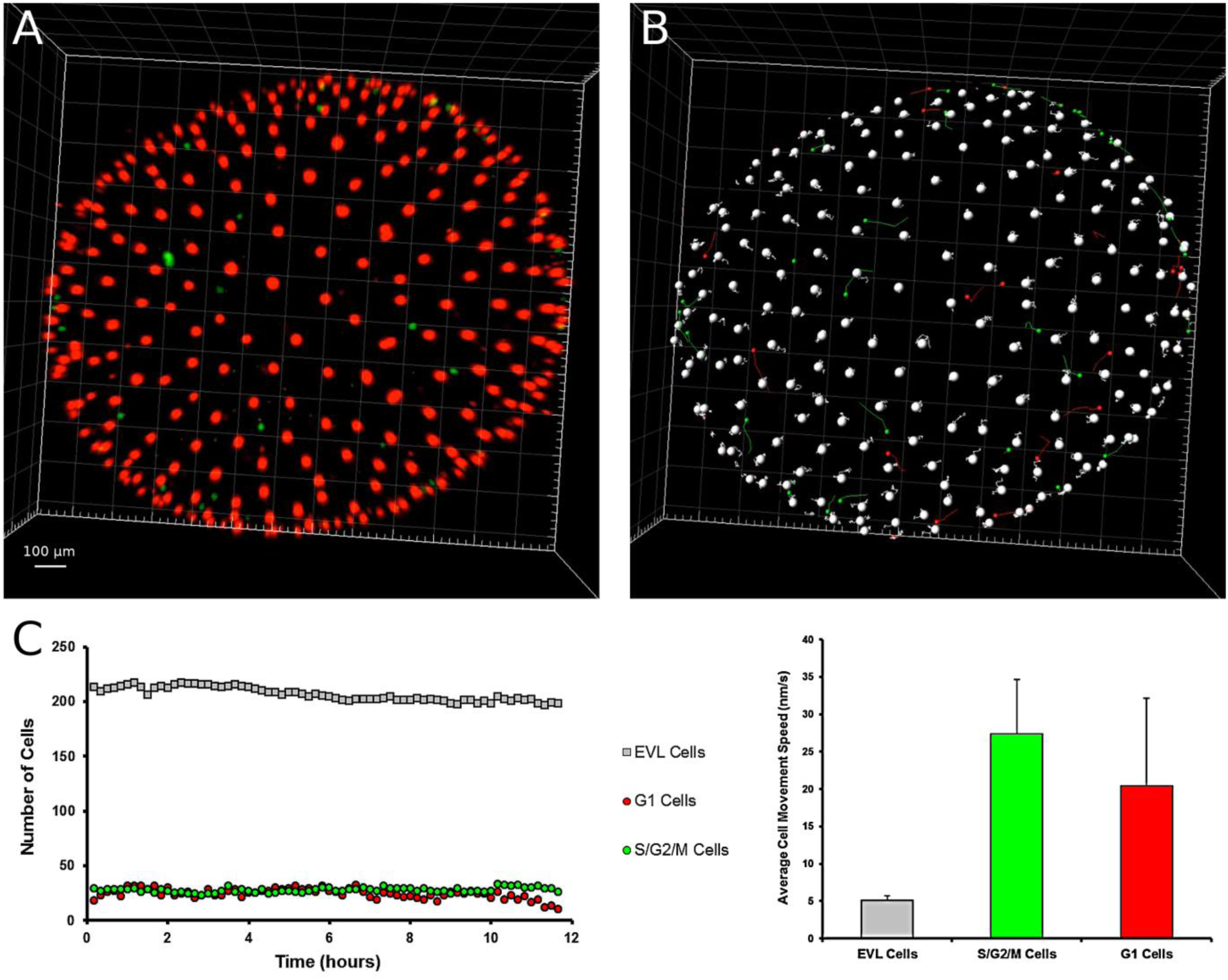
Cell dynamics during the early dispersed phase (Wourms stages 19-20). The first part of the dispersed phase is characterized by the presence of few EL green or red cells (A). ESL cells are pseudocolored as large white dots and EL green and red cells as small green and red dots, respectively (B). Tracking of the three cell types over time. The number of each type of cell was constant for more than 10 hours (C) and EL cells continuously moved during the whole time. Tracks of individual cells are shown in (B), and speed average values in C, right panel. The images and graph refers to the acquired portion of the embryos, equivalent to the superior hemisphere.

In addition, all the embryos we could observe at the end of epiboly (N > 30, including embryos that were not followed in time lapse but only through still imaging), transited through a phase when only few EL green or red cells were detectable, characterized by the following features:

- Regularly spaced EVL large cells nuclei (19-25 μm nucleus diameter) that do not move, migrate or divide (Figure 6).
- Presence of few (less than 80) red and green small epiblast cells (7-13 μm nuclear diameter) that do not divide (Figure 6).
- Seemingly random cell movements, along with “dark” cells visible only in brightfield (Movie as additional file 2).

Early dispersal occurred in all embryos and was reached by a continuous decrease of red and green fluorescent EL cells during epiboly (Figure 4). The duration of early dispersal is variable (from hours to days), likely depending on environmental conditions. This stage corresponds to diapause 1 (DI). Noteworthy, DI is characterized by a complete stop of the cell cycle with cell synchronized at the end of M phase. However, DI is far from being a static stage, as cells constantly move over the yolk surface (Movie as additional file 2, min 0.53 to 1.10). Therefore, DI may not necessarily have lower metabolic demands than normal development. The functional relevance of erratic cell movements remain, for the moment, unexplored.

### Synchronized cell-cycle re-entry underlies release from diapause I

The mechanisms responsible for release from diapause I are unknown, although previous experiments suggest that it is a temperature-dependent process [14, 47, 48]. The majority of embryos imaged during dispersed phase were either in a phase with few green cells (80 or less) or in a phase with a larger number of green cells (300 or more), indicating that the duration of the transition is considerably shorter than either of these two phases and difficult to capture. We were nonetheless able to image the exit from DI in four embryos. In all four cases, the release from DI was characterized by the rapid appearance of a large number of green fluorescent cells that was not preceded by a rise in red cells, indicating that: i) the “invisible” EL cells were indeed synchronized and ii) most of the “invisible” EL cells entered directly into S/G_2_/M phase of the cell cycle without transiting through G_1_ (Figure 7). Remarkably, these reactivated cells showed what seemed to be a pattern of synchronous division, generating a peak of green cells followed by a period of reduced cell numbers (likely because some cells entered in the “dark” phase). In two out of the four imaged embryos, subsequent pulses with a periodicity of 10 hours were apparent. Cells alternated from a green phase to a dark phase without showing a red phase, a pattern typical of early cleavage in *N. furzeri* [7]. Peaks of green fluorescent EL cells were present, but less apparent, in a third embryo whereas only the first peak was clearly visible in the fourth embryo (Additional file 3). This synchronization may simply reflect the fact that cells are synchronized during G1 and also cell cycle exit is synchronized. This is however a totally unique feature of embryonic development as cell cycle synchronization is otherwise known only for the very early phases of development that are controlled by maternal transcripts.

**Figure 7.**
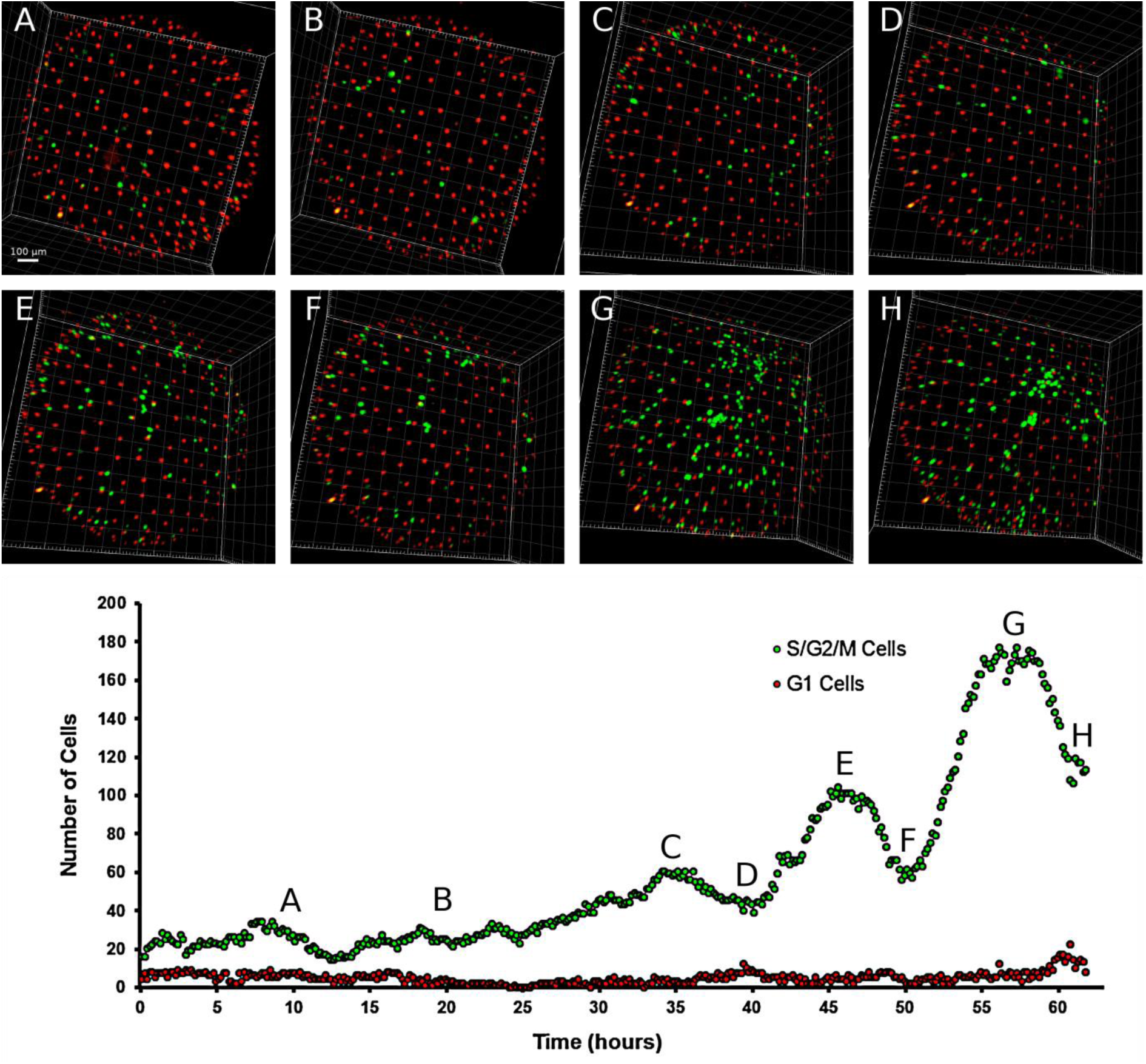
Transition from early to late dispersed phase. (A-H) Time lapse showing the transition from a stage with few EL red or green cells could be detected through multiple reactivation and division events up to a relatively stable condition where the number of green EL cells is ∼5 times their original number. (G) Quantification of green EL cell numbers over time. The letters A-H correspond to the pictures shown in (A-H). Multiple peaks of synchronized proliferation are clearly visible (C,E,G) and divided by phases when cells synchronously enter into G1 phase (D,F,H). The images and graph refer to the acquired portion of the embryos, equivalent to the superior hemisphere.

In all four cases, the number of cells increased over time-indicating active division- and the stage with ∼ 300 green cells was reached in about 36 hours. In addition, the proliferating cells moved erratically during the reactivation, in a way comparable to the movements of epiblast cells that were documented during DI by brightfield microscopy (Movieas additional file 2) or during early dispersed phase by confocal microscopy (Figure 4).

### Late dispersed phase (Wourms stage 20)

The late dispersed phase (Figure 8) was characterized by an almost equal number (∼ 500) of EVL, EL green and EL red cells. Random movements of EL cells continued at a speed comparable to that observed during the early dispersed phase and diapause I (Figure 6B and 5C, right panel). The detectable EL cells did not increase in numbers and were probably locked in G_1_ or G_2_. The amount of time embryos spent in this second part of the dispersed phase could not be determined. At the moment, we cannot offer an explanation for this remarkable phenomenon.

**Figure 8.**
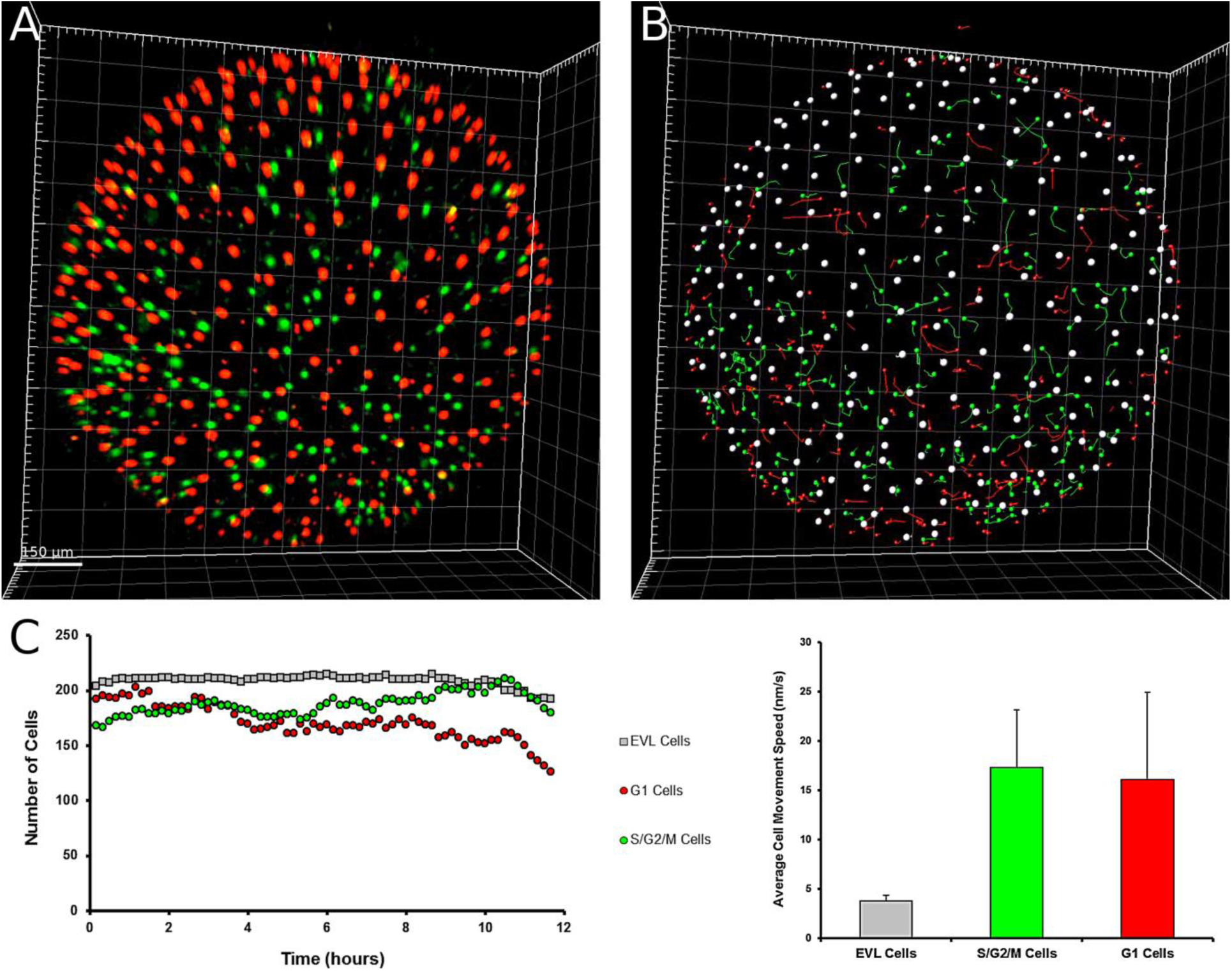
Cell dynamics during late dispersed phase (Wourms stage 20). The second part of the dispersed phase is characterized by the presence of a larger number of EL green or red cells as opposed to early dispersed phasAe (Figure4). (A-B) Cell tracking. EVL cells are visualized as large white dots while EL green and red cells as small green and red dots, respectively. All detected cells were tracked over time. (C) Quantification of the he numbers of each type of cell over time. Note that then number is relatively constant for more than 10 hours, while still moving. The images and graph refer to the acquired portion of the embryos, equivalent to the top superior hemisphere.

### Reaggregation (Wourms stages 21-26)

Reaggregation starts when the majority of EL cells change their movements from erratic to directed towards a specific region of the embryo, forming an initially sparse circular aggregate, whose radius becomes progressively smaller (Figure 9 and Movie as additional file 4 min 0.00 to 0.12). It was not possible to determine the point of the embryo toward which cells migrated in our experiments, but experiments performed by Wourms in 1972 [6] suggest that this is located in the lower hemisphere of the egg. The movements of the cells in this region were greatly reduced and the circular formation reduced its diameter over time. The reaggregation process required a small fraction of total developmental time and in about 15 hours the final circular aggregate of green cells was completely defined. A more extensive description of the aggregation and gastrulation processes in annual killifish and the associated cellular movements was recently provided by Pereiro et al. [9] and is beyond the scope of the present study.

**Figure 9.**
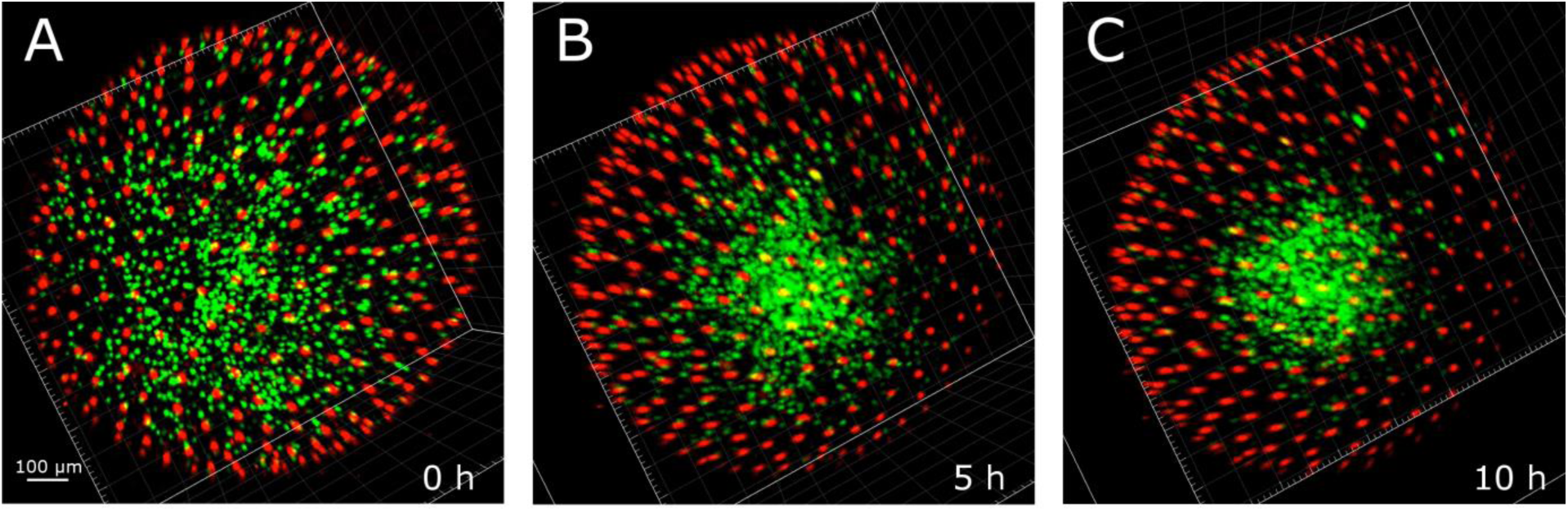
Reaggregation phase (Wourms stages 21-25). Green EL cells converge initially onto a single region of the embryo (A,B,C), forming a circular structure that over time becomes more compact with a reduced radius.

The circular organization progressively changes in shape without varying its area, becoming an ellipsoid that extends along its main axis retaining a higher density of green cells in the inner part and a decreasing density on the borders (Figure 10 and Movie as additional file 4min 0.12 to 0.21). Of note, the embryo is formed mainly by green cells with very little fraction of red cells, possibly because cell multiplication is necessary to reach a sufficient cell number for all the different embryonic structures to be formed and so the predominant cycle phases are the proliferation phase S/G_2_/M. In this phase, the number of epiblast green or red cells that do not belong to the embryo primordium and shows erratic movements is greatly reduced and is comparable or smaller than the number of cells that randomly move during diapause I (<80).

**Figure 10.**
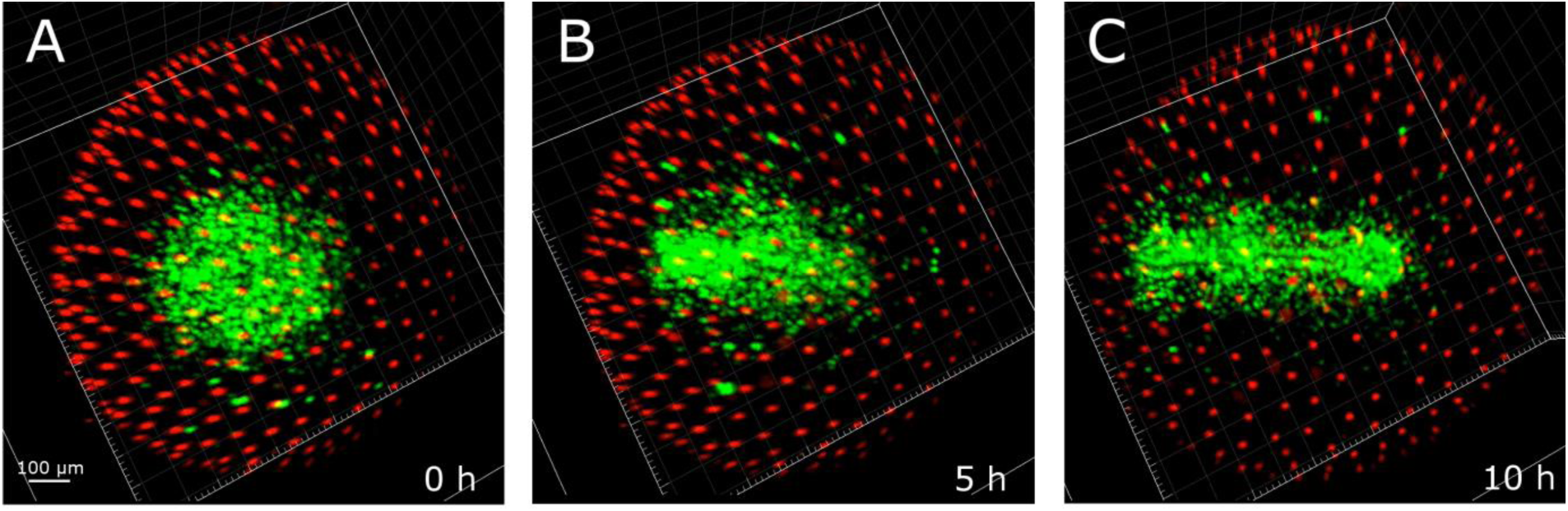
Axis extension phase (Wourms stage 26). The initial circular formation (A) lengthens to form the primordial axis (B-C).

### Somitogenesis (Wourms stages 29-33+)

The description of somitogenesis refers to direct developing (DD) embryos. We were unable to image any diapause committed (DC) embryo during this phase, most likely because the temperatures reached in the imaging chamber (> 26 °C) prevented diapause induction. Somitogenesis is associated to changes in expression of the FUCCI reporters that are highly reminiscent of those first described during zebrafish development [40]. The somites are the first structures where red fluorescence predominates. Their formation is progressive and is completed within a few hours, (Movie as additional file 4 min 0.26 to 0.52). Somites increase in number by addition of progressively more caudal somite pairs, as typical for teleost embryos (Figure 11).

**Figure 11.**
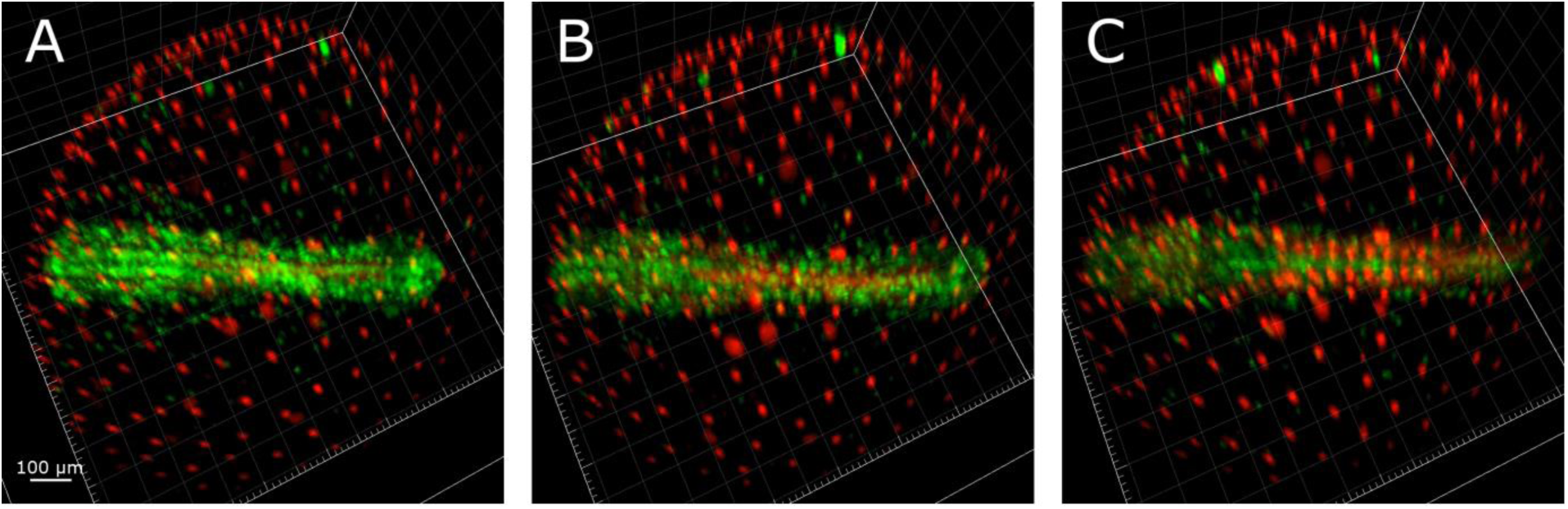
Direct developing embryos late somitogenesis (Wourms stages 29-32). Green cells populate the whole embryonic axis for all the somitogenesis in embryos that do not enter diapause II. Green cells number and density slowly drops as long as development proceed while red cells increase with time populating at first the somite pairs and afterwards almost every region of the axis.

After the formation of the first 2 somites, 4 symmetrical green streaks of proliferating cells become apparent, reaching from the caudal margin of the head to the end of the tail: two inner proliferation streaks, close to the midline and divided by the midline itself, and two outer streaks, defining the outer borders of the embryo (Additional file 5). The somite structures are located between the inner and outer streaks of green cells.

The two inner streaks contain cells that migrate inwards (Movie S5), while the movements of the cells of the outer streaks do not follow a clear direction. Some cells belonging to the outer streaks migrate outwards.

As somitogenesis proceeds, the green signal slowly reduces its intensity due the reduction in the fraction of proliferating cells and the expansion of red cells from the caudal somites in the anterior direction. However, even at late stages of somitogenesis, DD embryos are never exclusively dominated by red cells, which would be expected in case of a complete cessation of proliferation events, as for embryos in DII (Figure 13).

**Figure 12.**
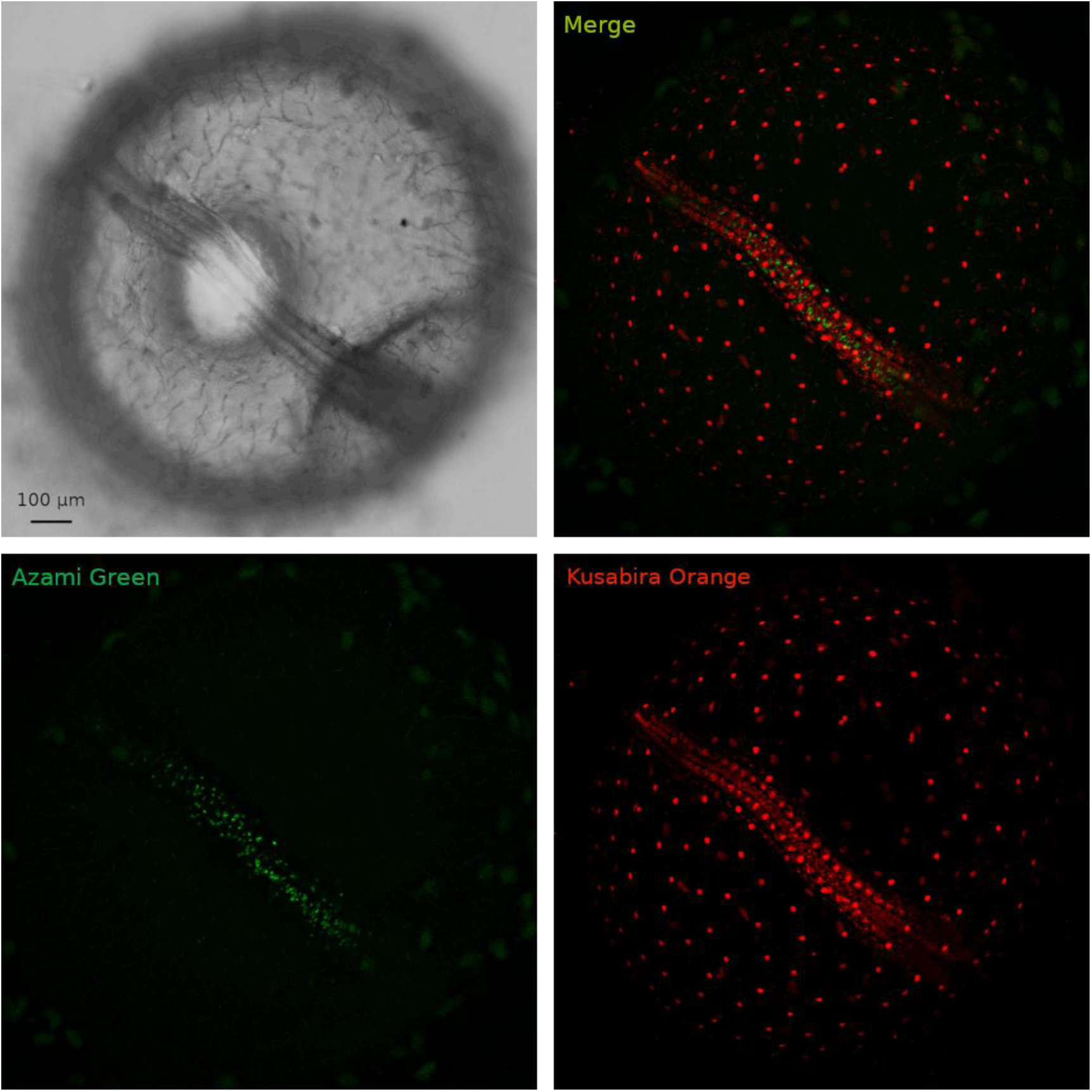
Diapause II arrested embryo. An embryos arrested in diapause II is illustrated. (B) Green cells are concentrated in the medial part of the embryo between the somites. (C) Red cells clearly define the already formed somites and diffusely populate the whole axis region. (D) overlay of green and red signal.

**Figure 13.**
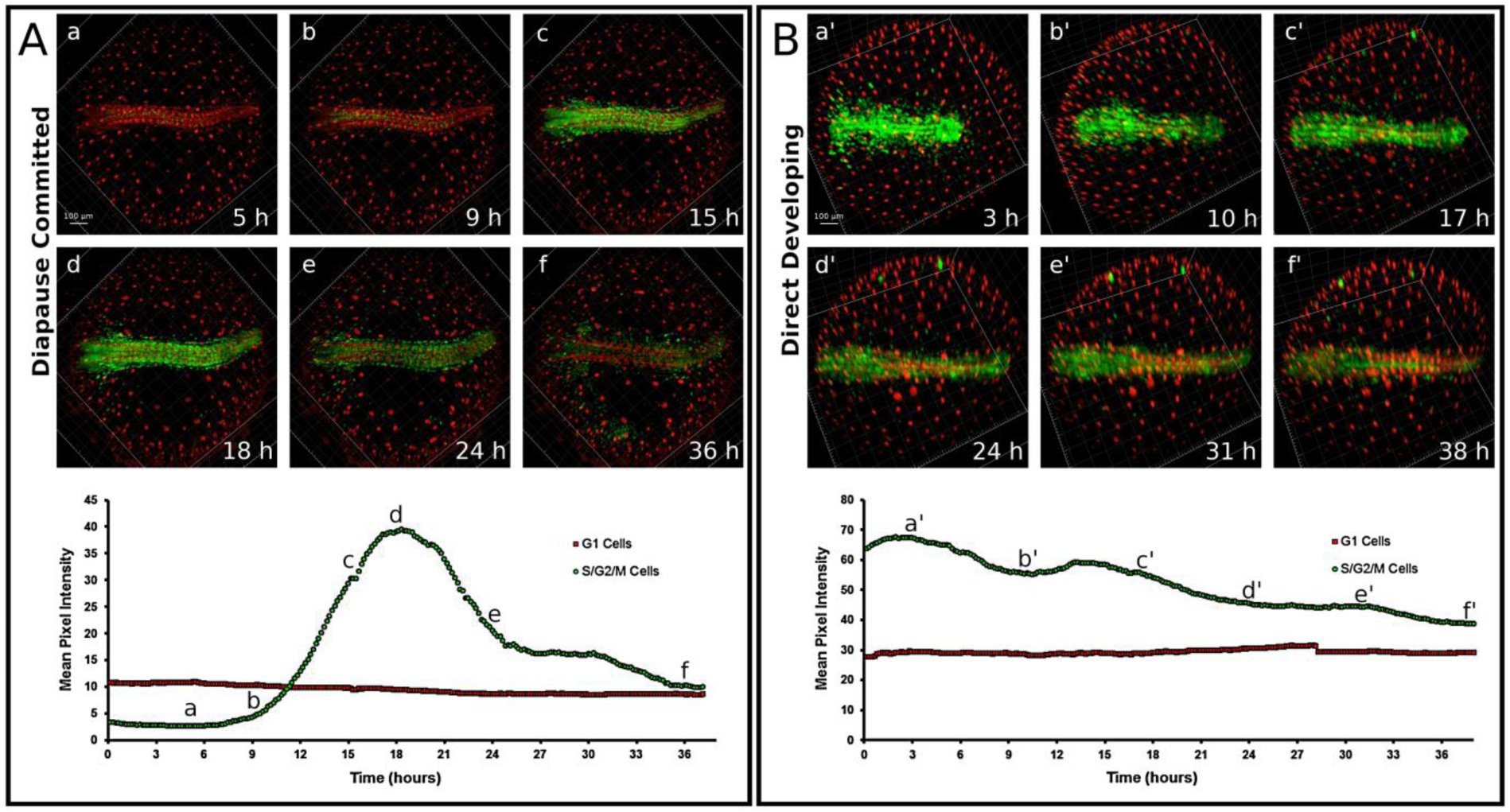
Timelapse of red and green fluorescence. Release from diapause II(A) as opposed to direct development (B). Upon release from diapause II, green cells greatly increase their number and density in less than 10 hours (A). After the initial proliferation burst (c,d), green cells slowly decrease in number and density (e-f). In the lower panel, a quantification of green signal intensity as an indirect estimate of the number of green cells is reported. The lettering indicates the correspondence of the curve with the pictures (a-f). (B) An example of direct development. The number of green cells decreases gradually while the number of red cells is roughly constant (a’-f’). Note that at the end of the processes the direct developed embryo (f’) is very similar to the embryo that escaped diapause (f). In the lower panel, a quantification of green signal intensity as an indirect estimate of the number of green cells is reported. The lettering indicates the correspondence of the curve with the pictures (a’-f’). Note that at the end of the processes, the ratio between red and green fluorescence in the direct developed embryo (f’ in the right panel) is similar to that of the embryo that escaped diapause (f in the left panel).

### Diapause II

In order to obtain images from embryos in DII, these were incubated for weeks at low temperature (22 C°) before imaging. At the beginning of the imaging, diapausing embryos contained almost exclusively cells with red fluorescence, indicating an almost complete suppression of proliferation. Sparse green cells could be detected along the midline and spread between the somites, but their number was greatly reduced compared to DD embryos. In addition, these remaining green cells were possibly blocked in G_2_ and not dividing, since the green signal did not increase (Figure 12). For the technical reasons delineated above, it was not possible to document diapause entry and therefore it remains unclear whether cell cycle arrest in embryos committed to diapause (DC) is progressive or it is reached abruptly at a specific developmental stage.

### Synchronous cell cycle re-entry and catch-up growth upon release from diapause II

The release from diapause II was documented in a total of 4 embryos (Movies as additional files 6 and 7) and in all cases it was characterized by a rapid catch-up process by which almost all the previously red or colorless cells of the embryo, with the exclusion of the cells forming the somites, switched to green fluorescence, showing a reactivation of proliferation (Figure 13A and Movie as additional file 6). Cell cycle reactivation starts apparently simultaneously in the whole embryo without an antero-posterior gradient and is a rapid process with green signal doubling in less than four hours and reaching a peak in about ten hours (Figure 13A b-d, additional file 8). As in the case of release from DI, the upregulation of green fluorescence is not followed by an increase of red fluorescence, suggesting that cells proceed through S/G_2_/M phases without long permanence in G_1_. This burst is transient, once the peak is reached, the green signal halves within about five hours and reaches a steady state within one day (Figure 13A, d-f, additional file 8). Our analysis confirms that DC and DD are distinct developmental pathways. Indeed, green signal is reduced but always present in DD embryos, while diapausing embryos show almost total absence of green cells. Another difference between DC and DD is the smaller axis thickness of DC embryos that was already described by Furness et al, [15]. The reactivation results primarily in a widening of the embryo with little longitudinal growth. Indeed, quantification of the inter-somite distance showed a steady increase during this phase (Figure 14).

**Figure 14:**
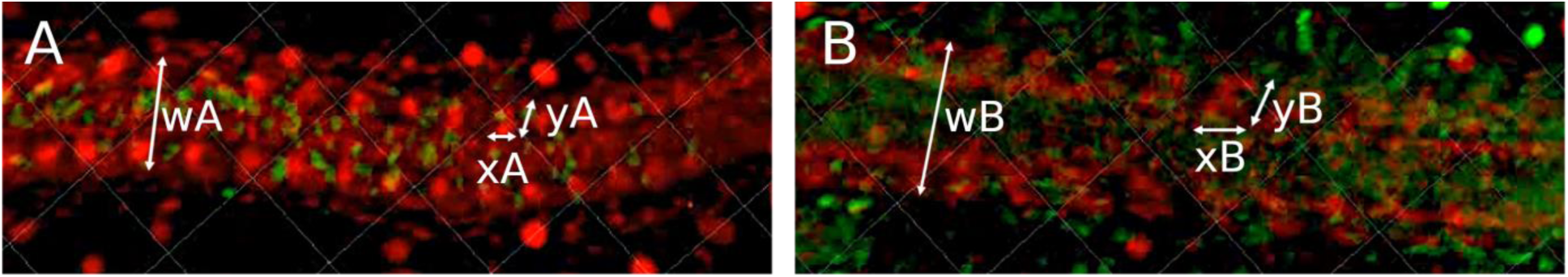
Change in somite morphology during release from diapause II. (A) start and (B) end of the catch-up phase. Reported are axial width (w), somite width (x) and somite length (y). wA=70 ± 6 um; wB= 97 ± 9 um; xA= 16.5 ± 2.5 um; xB= 24.6 ± 2.8 um; yA= 16.7 ± 2.4 um; yB= 25.5 ± 3,5 um. Measurements performed with the software FIJI.

Reactivation causes the somites to change morphology (Figure 14). The somites of a DII embryo have a roughly circular morphology. At the end of the reactivation period, their shape is elongated, at an angle of roughly 45° with the embryonic axis. Therefore, within less than one day, a DC embryo changes its morphology to become indistinguishable from a DD embryo. The rapid change in morphology in somites and embryos upon reactivation is compatible with the observation that embryos in diapause upregulate genes responsible for protein synthesis, such as ribosomal proteins and initiation/elongation factors [36], despite protein synthesis being suppressed in diapausing embryos [34]. Diapausing embryos could be “primed” to rapidly recover growth, as translation could be more rapid and efficient in re-entering the cell cycle and supporting the metabolic needs of a growing embryo.

## Conclusions

Our study improves the existing knowledge on annual killifish embryology and defines the following different phases of *N. furzeri* development based on cell cycle properties that are summarized in Figure 15. Of particular interest are the following points:

- An early dispersed phase is characterized by cells possibly locked at the end of the M phase;
- Diapause I is a prolongation of the early dispersed phase when cells randomly move but do not divide;
- Exit from diapause I is characterized by synchronized bursts of cell division;
- After exit from diapause I, embryos do not proceed immediately to reaggregation, but transit through a late dispersed phase with random migration of epiblast cells without cell division
- Exit from diapause II is characterized by a transient burst of cell division and a catch-up process that leads primarily to transversal growth of the embryo with little longitudinal growth.

**Figure 15.**
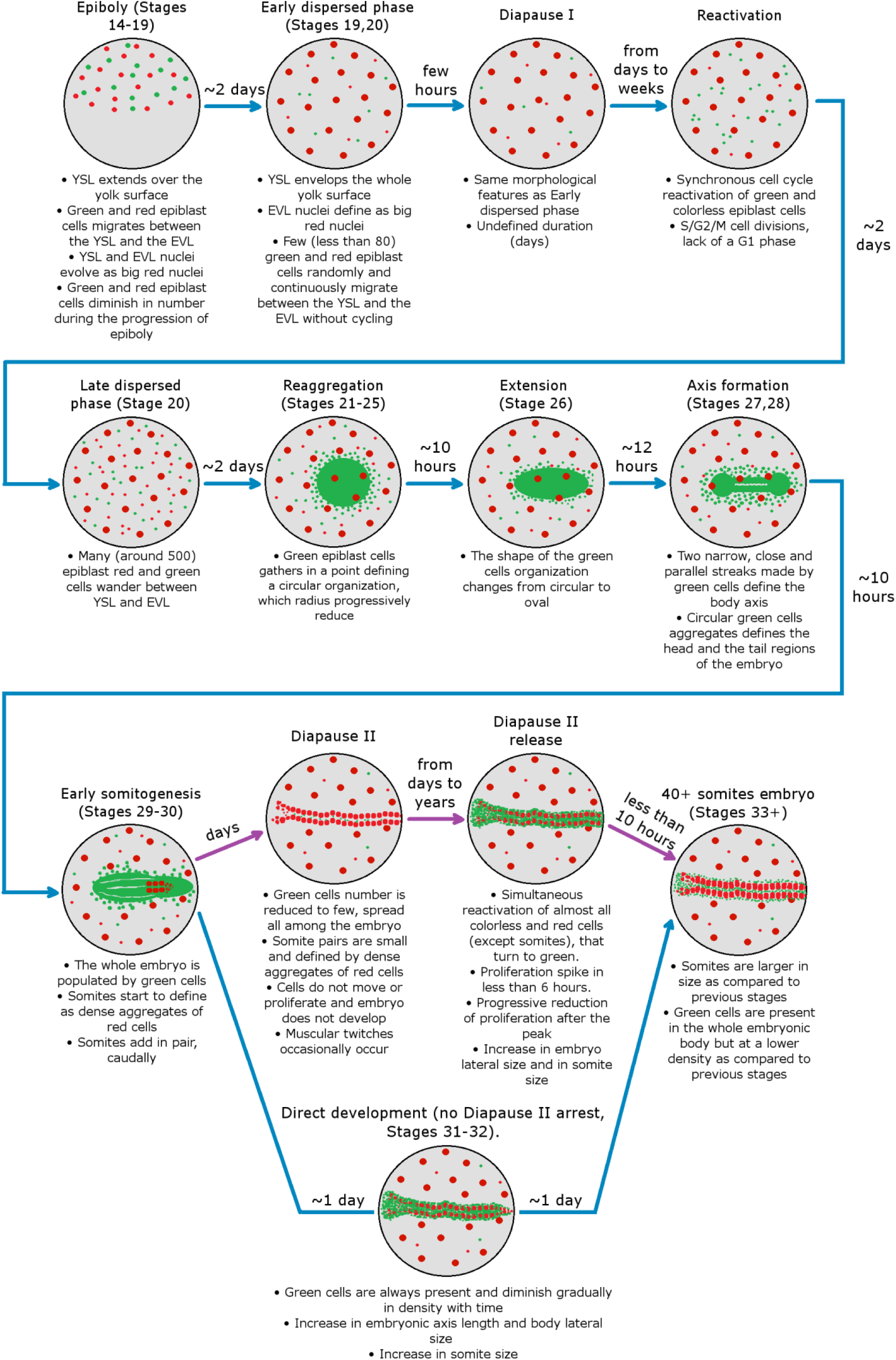
Graphical abstract representing FUCCI N. furzeri fish development. Images and developmental times refers to 26°C incubation conditions, except purple arrows that refers to low temperature (18°C-21°C) incubation conditions. Stages refers to developmental stages described by Wourms [6].

In addition to the new knowledge we have derived on cell dynamics during embryonic development, we imagine that the FUCCI transgenic line may represent a resource with several applications.

In a first instance, it allowes a very precise identification of the timepoint when an embryo exits from DII.

Release from DII is not synchronous, possibly as a strategy of bet hedging [10]. Currently, embryos that exit DII can be only identified based on embryo width, eye size or pigmentation by daily observations. This technique can reliably identify only embryos that left DII a few days in the past and became apparently different from diapausing embryos. With our tool, the time window between DII exit and expression of a detectable signal is narrowed to few hours. This transgenic line, combined with FACS sorting and transcriptomics/proteomics, could offer a new and unique entry point to dissect the molecular mechanisms responsible for exit from quiescence.

A second perspective, that goes beyond developmental dynamics, is the study of cell dynamics in adult organs. In particular, *N. furzeri* represents a convenient model to study the effect of aging on adult stem cells [49] and tail regeneration [50]. FUCCI offers a direct tool to quantify changes of cell cycle dynamics during aging and in response to regeneration and to purify and analyze adult stem cells/progenitors.

Finally, *N. furzeri* is characterized by a high incidence of age-dependent spontaneous tumors. FUCCI would provide a tool to visualized and investigate tumors in vivo by intravital microscopy [51], allowing longitudinal studies of spontaneous tumorigenesis in living fish, characterization of tumor response to pharmacological treatments or other interventions and sorting of tumor-derived cells for molecular analysis.

## Methods

### Declarations

#### Ethics approval

The experiments described do not classify as “animal experimentation” under the Directive 2010/63/EU as the transgenic lines used do not have a suffering phenotype. A general licence for animal housing, breeding and manipulations was issued by the Umwelt- und Verbraucherschutzamt der Stadt Köln. Authorization Nr. 576.1.36.6.G28/13 Be.

## Data availability

All data generated or analysed during this study are included in the supplementary information files or can be obtained by the corresponding author on a reasonable request.

## Authors contributions

L.D. and A.C. designed the study, L.D. and R.R. created the transgenic lines, L.D. performed the imaging and data analysis. A.A., D.R.V. and A.C. supervised the study. L.D. and A.C drafted a first version of the manuscript. L.D., A.A., D.R.V. and A.C. wrote the final version of the manuscript.

### Acknowledgements

The authors thank X,Y,Z for technical assistance and Eva Terzibasi Tozzini for help in fish husbandry during an initial phase of the project.

## List of additional files

Additional file 1: Movie of particle tracking during epiboly

Additional file 2: Bright-field time-lapse of early dispersed phase

Additional file 3: Graph depicting the changes of fluorescence in four embryos released from diapause I.

Additional file 4: Time-lapse fluorescent imaging of reaggregation phase, axis formation and early somitogenesis

Additional file 5: Still fluorescent image of a direct-developing embryo

Additional file 6: Time-lapse fluorescent imaging of exit from DII

Additional file 7: Time lapse, comparison of four different embryos that exit from DII

Additional file 8: Quantification of the fluorescence of the four different embryos shown in additional file 7.

